# Analytical Ultracentrifugation as a Tool for Exploring COSAN Assemblies

**DOI:** 10.1101/2025.01.30.635703

**Authors:** Hussein Fakhouri, Caroline Mas, Aline Le Roy, Estelle Marchal, Coralie Pasquier, Olivier Diat, Pierre Bauduin, Christine Ebel

## Abstract

The self-assembly of the cobalta bis(dicarbollide) (COSAN) anionic boron clusters into micelles above a critical micelle concentration (cmc) of 10 - 20 mM and its behavior as “sticky nano-ions” facilitating controlled protein aggregation have been previously investigated using scattering techniques, particularly small-angle neutron and X-ray scattering. These techniques effectively provide average structural parameters but, when applied to colloidal systems, often rely on models assuming polydispersity or anisotropic shapes. Thus, complementary techniques are required to confirm or infirm the proposed analyses. Here, we employed sedimentation velocity analytical ultracentrifugation (SV-AUC), which offers the ability to resolve discrete species. We revisited two key questions: (1) the aggregation behavior of COSAN into micelles, a topic still under debate and due to its unconventional amphiphilic nature, and (2) the nature of the protein assemblies induced by COSAN, specifically their size/shape distribution and aggregation number. Our findings confirm the cmc of COSAN of 16 mM and reveal that COSAN micelles exhibit low aggregation numbers (8 in water and 14 in dilute salt), consistent with recent hypotheses. Furthermore, SV-AUC showed that COSAN promotes myoglobin aggregation into discrete oligomeric species with well-defined aggregation numbers, such as dimers, tetramers, and higher-order assemblies, depending on the COSAN-to-protein ratio. These results provide clarity on the discrete nature of COSAN micelle aggregation and protein assembly. This study highlights the complementary role of SV-AUC in understanding supramolecular assemblies, offering useful insights into the behavior of COSAN nano-ions and their interactions with biomacromolecules.

## Introduction

Cobaltabisdicarbollide (COSAN), [3,3′-Co(1,2-C_2_B_9_H_11_)_2_]^−^, is a boron-based anion featuring a cobalt(III) cation sandwiched between two hydrophobic dicarbollide semicages and carrying a formal -1 charge (Figure 1) (Grimes 2016). This charge is delocalized over the nanometric structure, resulting in a low charge density anion with properties characteristic of a superchaotropic species (Assaf et al. 2015; Naskar et al. 2015). The superchaotropic effect of COSAN has been specifically studied in previous research (Assaf et al. 2019; Merhi et al. 2020). The surface of COSAN exhibits weakly polarized B-H and C-H bonds, conferring a hydrophobic character and the propensity to form non-specific H-bounds with its environment. This combination grants COSAN an intrinsic hydrophilic/hydrophobic duality. Despite lacking the classical hydrophilic-hydrophobic segmentation typical of surfactant molecules, COSAN exhibits all the hallmark properties of surfactants, including surface activity (Popov and Borisova 2001; Gassin et al. 2015), foaming (Viñas et al. 2014), self-assembly (Bauduin et al. 2011; Matějíček et al. 2006), and the formation of lyotropic lamellar phases at high concentrations (Brusselle et al. 2013).

**Fig. 1.**
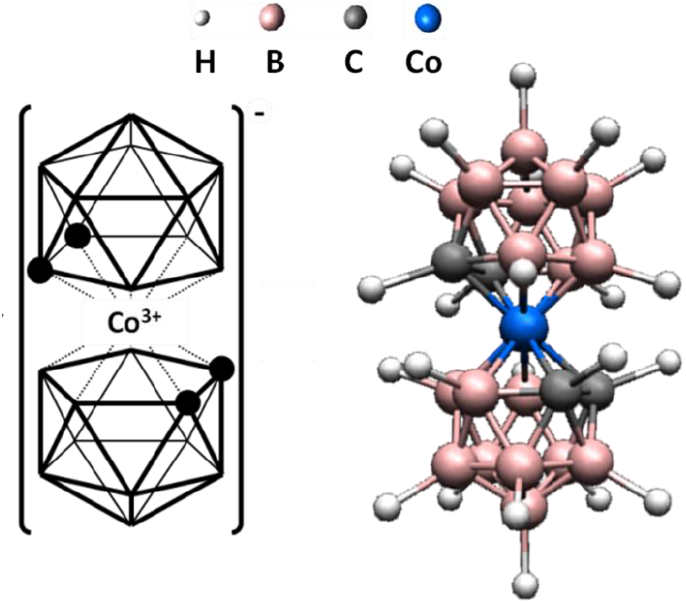
COSAN Structure.

COSAN’s self-assembly in water is particularly unique: at low concentrations, it forms large monolayered vesicles, which transition into small micelles at higher concentrations, both assemblies in equilibrium with COSAN monomers (Bauduin et al. 2011). This behavior contrasts with classical organic surfactants, which typically form either micelles or vesicles, depending on their critical packing parameter (Israelachvili 2011). It has been proposed that, upon increasing concentration, charged COSAN vesicles undergo a Coulombic explosion, causing the vesicle to–micelle transition. Small-angle neutron scattering (SANS) on COSAN micellar solutions produces a scattering signal that can be well-fitted using a sphere model with electrostatic interactions. The resulting radius corresponds approximately to the long axis of the COSAN anion. SANS experiments suggested a critical micelle concentration (cmc) of 18.6 mM (Bauduin et al. 2011), whereas NMR and surface tension measurements yield a lower cmc value closer to 10 mM (Uchman et al. 2015). Based on the micelle size obtained from SANS data fitting, the maximum aggregation number has been estimated to be around 15 when assuming the sphere’s volume is entirely filled with COSAN molecules, and approximately 12 when considering molecular packing constraints. Recently, building on previous investigations, a study carried out by Medož and Matejíček (Medoš et al. 2022) revisited COSAN aggregation in water using isothermal calorimetry and supported by molecular dynamics (MD) simulations. They proposed that COSAN assembles initially as pentamers, followed by a second-step aggregation process at higher concentrations, leading to aggregates with an average aggregation number of 9 ± 2. Furthermore, they highlighted the possible critical role of counterions (sodium) in inducing and stabilizing COSAN self-assembly. MD simulations revealed a highly polydisperse aggregation pattern, with species ranging from monomers to dodecamers, while pentamers and hexamers were the predominant species at 50 mM COSAN.

Beyond its remarkable self-assembly behavior, COSAN also exhibits a strong capacity to interact with other molecules, opening avenues for diverse biological and therapeutic applications (Gabel 2015; Leśnikowski 2016). Its exceptional stability, arising from 3D aromaticity (King 2001), combined with high water solubility, biocompatibility, chemical inertness, pronounced chaotropic and hydrophobic characters (Merhi et al. 2020; Chen et al. 2023), makes COSAN a versatile scaffold in medicinal chemistry (Fuentes et al. 2018). These properties have enabled its use in boron neutron capture therapy (Palmieri et al. 2024), radioimaging (Hawthorne and Maderna 1999), and magnetic resonance imaging, as well as in the development of enzyme inhibitors (Grüner et al. 2019) and antimicrobial agents (Kubiński et al. 2022). Furthermore, COSAN’s chaotropic nature facilitates the intracellular delivery of hydrophilic molecules, such as peptides, which are otherwise impermeable to cell membranes (Chen et al. 2023). Other ionic boron clusters with hydrophobic and chaotropic characteristics, such as the iodinated dodecaborate (B_12_I_12_^2-^), were recently found to accumulate in the phosphatidylcholine polar head regions of lipid bilayers (Barba-Bon et al. 2024). This association leads to a thinning of the membrane, which facilitates the transmembrane transport of hydrophilic molecules. COSAN is also known to bind to proteins (Goszczyński et al. 2017; Fuentes et al. 2019; Chazapi et al. 2024), exhibiting specific interactions, such as with HIV protease (Cígler et al. 2005), as well as non-specific binding with various proteins like myoglobin, carbonic anhydrase, and trypsin inhibitor. Recently, COSAN and B_12_I_12_^2-^ were additionally found to act as reversible molecular glue for proteins, inducing the formation of 2D assemblies with these proteins while preserving their secondary structures and even preventing thermal denaturation in the case of myoglobin. These interactions highlight the importance of understanding its protein-binding mechanisms to fully exploit its potential in drug delivery, gene therapy, and other biological applications. The size, shape, and aggregation number of these COSAN-protein assemblies were determined using small-angle X-ray scattering (SAXS), employing a three-axial geometrical model through spatial averaging, as is typical for all scattering techniques. While SAXS is well-suited for providing averaged spatial information about a population of species, it has significant limitations in determining precisely the aggregation number in these anisotropic aggregates, particularly when size differences are small. In the present case, fitting the data with a monodisperse anisotropic shape model can produce a scattering pattern indistinguishable from that obtained using a polydisperse model with a fixed shape. This highlights a fundamental limitation of SAXS, making it poorly adapted to differentiate between these scenarios. To overcome these limitations, other techniques such as cryo-electron microscopy and/or sedimentation velocity analytical ultracentrifugation (SV-AUC) may be used.

SV-AUC is indeed a powerful biophysical technique to determine the size, shape and aggregation state of macromolecules in solution and study their interactions. This method is widely applied in the characterization of both biological (Schuck 2013; Demeler 2024) and colloid samples (Cölfen 2023). SV-AUC experiments observe the movement of macromolecules in solution when submitted to centrifugal force. An optical system allows to measure sedimentation velocity (SV) profiles. SV profiles are concentration distributions, determined by *e. g*. absorbance or interference fringe shifts, measured as a function of the distance to the rotation axis, at different times of centrifugation. Each SV profile reflects, at a given time, the radial concentration distributions for the one or several types of macromolecules present in solution. Before spinning, and as observed typically at low angular velocity (600 *g*), the solution is homogeneous, and the SV profile is flat. As the solution is spun at high centrifugal field, (typically 130 000 *g*), macromolecules move down the centrifugal field, and solution-solvent boundaries are generated and evidenced as a significant signal change, in a typically restricted radial range, in the SV profile. The position and shape of the boundaries change with time of sedimentation. The whole set of SV profiles -recorded over typically 15 hours-is interpreted by the analysis (see material and method section). SV-AUC combines the separation of the macromolecules under gravitational force with a rigorous analysis of their transportation providing insights into their hydrodynamic and thermodynamic properties. The materials and methods section provides a more detailed description of the technique, while the principles of the analyses are here briefly exposed. The sedimentation and diffusion coefficients, *s* and *D*, determine the sedimentation of the macromolecules; they are directly related to their molar mass and hydrodynamic diameter, *M* and *D*_H_. The *c*(*s*) analysis yields a high-resolution size distribution, *c*(*s*), of *s*, which is particularly useful for probing sample homogeneity and identifying equilibrium of association by comparing samples at different concentrations. The *s*-values can be translated into *M*-values –thus aggregation numbers-with hypothesis on the frictional ratio, *f*/*f*_0_, which links *M* and *D*_H_. This is possible because the value of *f*/*f*_0_, which depends on the macromolecule compactness, shape anisotropy and hydration, evolves only in a limited range. The non-interacting species (NIS) analysis, can be used in favorable cases, to determine *M* and *D*_H,_ without prior hypotheses about the shape of the macromolecule. However, this is possible when well-resolved, discrete sedimenting species are present or if reversible interactions are slow under the experimental conditions. In both analysis, an estimate of the value of the partial specific volume of the macromolecule, *v*, is required to derive *M*. Combining different optical detections, using interference and/or absorbance at one or more wavelengths, in SV-AUC allows for the determination of stoichiometry in multicomponent systems, *e. g*. solubilized membrane proteins or adeno-associated virus vectors composed of protein, and detergent or nucleic acid, respectively (Le Roy et al. 2015; Saleun et al. 2023; Henrickson et al. 2023).

We aim to use SV-AUC to further investigate COSAN’s self-assembly behavior by distinguishing between various aggregated forms of COSAN, from monomers to larger assemblies, such as pentamers, hexamers, or dodecamers, under different concentration and salt (buffer) conditions. By analyzing the sedimentation coefficients, we expect to extract valuable data on the size distribution and heterogeneity of these species, refining our understanding of the aggregation process and cmc. In addition, SV and AUC will be used to explore COSAN-myoglobin assemblies, allowing us to discriminate between different protein aggregation states and gain a deeper understanding of the interactions at play. These techniques will overcome the limitations of SAXS/SANS, particularly in distinguishing between discrete and polydisperse aggregates, and provide high-resolution size and shape information, complementing data from other methods such as SAXS and molecular dynamics simulations. This integrated approach will offer valuable insights into the structural dynamics of COSAN’s self-assembly process and COSAN-protein assemblies.

## Results

### COSAN in water and dilute salt buffer

Different concentrations of COSAN in the range 1 - 100 mM (0.35 - 35 mg mL^-1^), in water and in dilute salt buffer (20 mM Tris-HCl pH 8, 150 mM NaCl, or PBS pH 7.4), were analyzed in SV-AUC. We typically used absorbance at 445 nm, *A*_445_, and fringe shift from interference optics, Δ*J*, to analyze the data. COSAN has two absorbance maxima at 280 nm and 445 nm. The extinction coefficients are reported in Table 1. With the smallest available optical path length of 0.15 cm, SV-AUC allows to characterize COSAN at concentrations up to ≈ 0.35 mM at 280 nm and ≈ 25 mM at 445 nm. Interference optics allows measurements typically between ≈ 0.1 (0.3 mM) and dozens of mg mL^-1^. Limitation at high concentrations may arise and be eventually corrected (Chaturvedi and Schuck 2024), from optical distortion related to the large fringe shift gradient within the boundary. This limitation does not apply for our COSAN samples, which sediment forming very broad boundaries, as will be shown below. The figures 2a and 2b present the sedimentation velocity profiles obtained at 445 nm and using interference optics, respectively, of COSAN at 5 mM: COSAN sediments slowly (≈ 0.2 S), without forming a clear boundary. The *c*(*s*) analysis of the data obtained from both optical systems are nicely superposed (Figure 2c), as expected for a solution with a single type of macromolecule. No large objects, such as vesicles, reported in (Bauduin et al. 2011; Matějíček et al. 2006) are detected. If such large objects exists in the solution, they are not present in detectable amounts here, or they are large enough (> 10^6^ Da) to be pelleted before acquisition of the first SV profile. This latter hypothesis is plausible, as the aggregation number for such large objects (vesicles) was estimated using static light scattering to be around 12,500, corresponding to a molar mass of approximately 4 × 10^6^ Da (Bauduin et al. 2011).

**Table 1.**
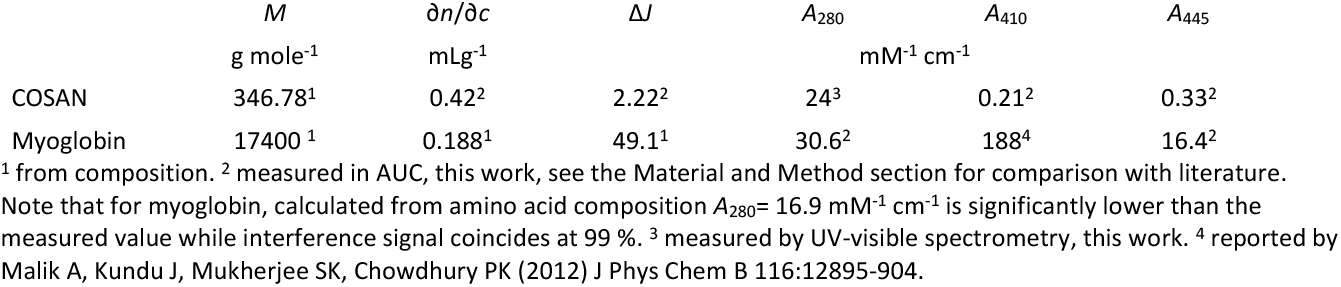
Extinction coefficients and ∂*n*/∂*c* values for COSAN and myoglobin.

**Fig. 2.**
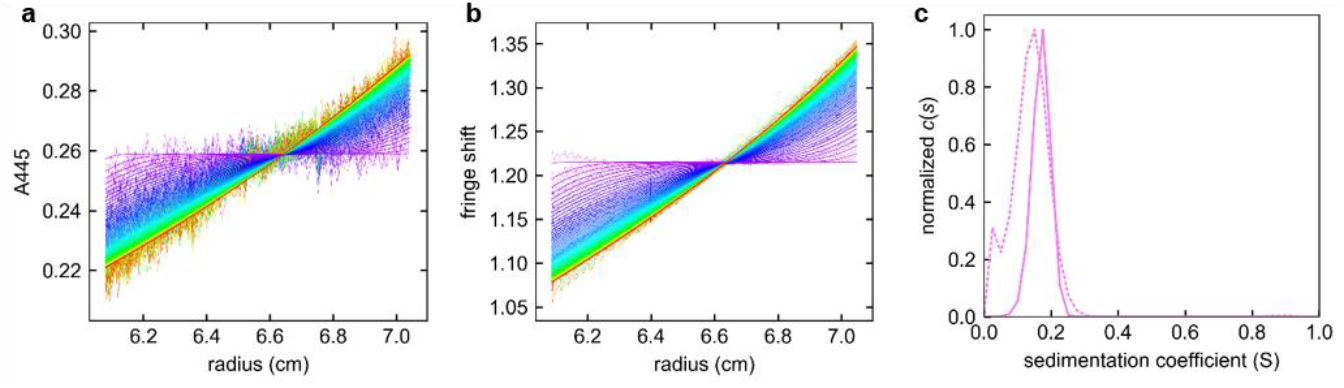
Sedimentation of 5 mM COSAN in H_2_O. Superposition of experimental and fitted sedimentation velocity profiles obtained for 5 mM COSAN in water. The sample was loaded in a 0.15 cm optical path centerpiece and centrifuged at 20°C at 42000 revs per min (130000 *g*) for 20 h. Data were acquired every 15 minutes at 445 nm (**a**), and using interference optics (**b**). The corresponding *c*(*s*) distributions plots obtained from *A*_445_ (continuous line) and interference optics (dotted line) are shown in (**c**).

Figure 3 presents the *c*(*s*) distributions measured at different COSAN concentrations from 1 to 100 mM in dilute salt buffer. They present a main peak with a mean *s*-value that increases from ≈ 0.1 S up to 0.7 S when increasing COSAN from 1 to 10 mM (Figure 3a). When further increasing COSAN concentration from 10 mM, to 100 mM, the *s*-value decreases (Figure 3b). The same trend is observed in water (Figure 4a): the mean *s*-value increases from ≈ 0.1 S to 0.45 S with COSAN up to 15 mM, and then decreases for COSAN above 15 mM. These changes suggests the auto-association of COSAN into small assemblies, the COSAN micelles, up to 10 or 15 mM and repulsive interactions between them above these concentrations. The mean *s*-value at the lowest concentrations of 1 and 2 mM COSAN, 0.11 +/- 0.04 S, coincides with the value of 0.13 S calculated for a globular monomer of COSAN, calculated with a frictional ratio of 1.25, and a partial specific volume of 0.736 mL g^-1^ (Medoš et al. 2022a). COSAN below 2 mM was indeed described to be in the monomer state (Medoš et al. 2022). COSAN at 5 and 10 mM sediments as a monomer in water at 0.15S, while in diluted salt solvent it sediments faster at ≈ 0.3 and 0.6S (see the Figure 4a). This suggests that the transition from monomer to micelle is decreased by around 5 mM for the dilute salt solvent when compared to water, in line with previous characterization (Fernandez-Alvarez et al. 2018). AUC-SV does not allow distinguishing whether the observed increase in the mean sedimentation coefficient corresponds to the presence of intermediate aggregates or results from a rapid equilibrium between monomers and well-defined micelles.

**Fig. 3.**
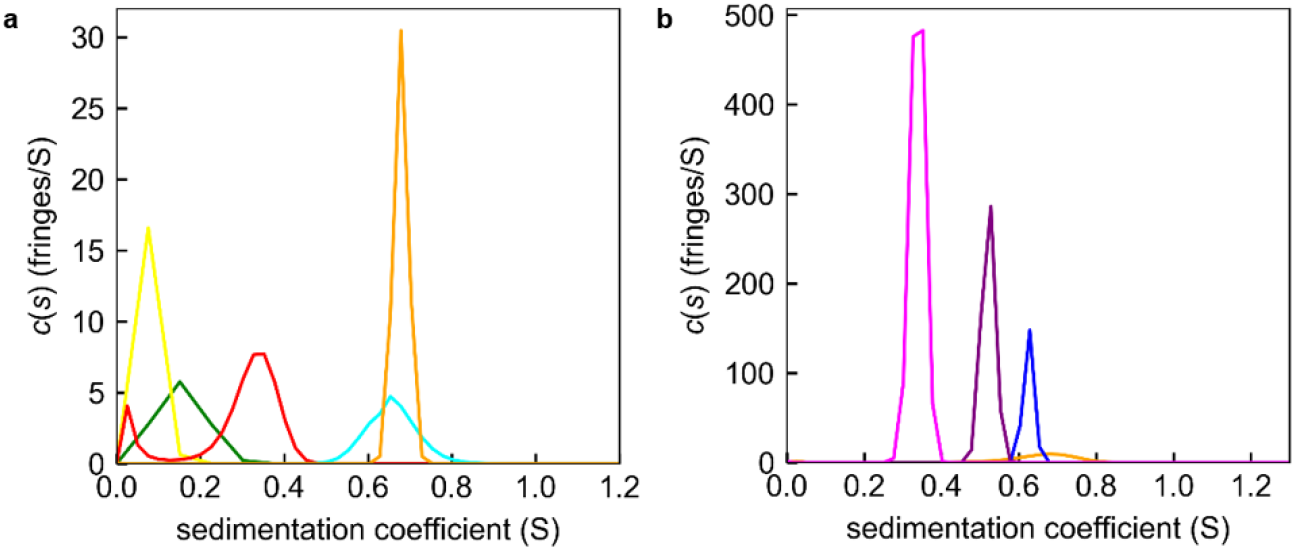
Sedimentation of COSAN up to 100 mM in water and dilute salt buffer. *c*(*s*) analysis of COSAN sedimentation followed by interference optics, **a**: at 1, 2, 5, 10, and 15 mM (green, yellow, red, cyan and orange respectively) in 20 mM Tris –HCl 150 mM NaCl pH 8; **b**: at 15, 30, 60, and 100 mM (orange, blue, purple and magenta respectively) in PBS pH 7.4.

**Fig. 4.**
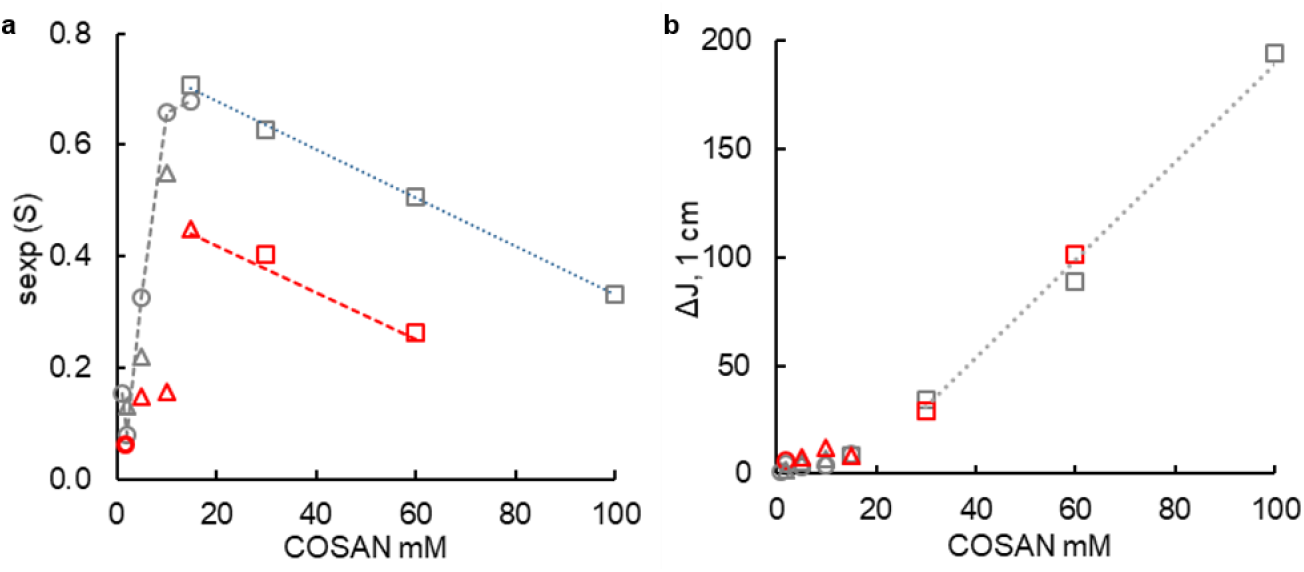
Sedimentation coefficients and interference signals measured in AUC for COSAN in water and dilute salt buffer. **a**: sedimentation coefficient **b:** Fringe shift displacement, Δ*J*, derived from the *c*(*s*) analysis, for COSAN in water (red square, triangle and circles, for different experiments), or in 0.15 M NaCl buffered with 20 mM Tris-HCl pH 8 (grey circles), or PBS (grey square and triangle, for two different experiments). Linear regressions were done, in (**a**), above 15 mM, to extrapolate the sedimentation coefficients at infinite dilution and the non-ideality hydrodynamic parameter *k*_s_, and, in (**b**), above 30 mM, to determine the cmc and ∂*n*/∂*c*. Linear regression considering all data in water and in dilute salt buffer, is Δ*J* _1 cm_ = 2.27 [COSAN] (mM) - 37.54, R^2^ = 0.9992.

The sedimentation coefficients, in water as in PBS, linearly decrease above 15 mM (5.2 mg/mL). It is very general that macromolecules sediment slower for concentrated solutions - above the mg/mL range-compared to diluted ideal solutions (Solovyova et al. 2001; Chaturvedi et al. 2018).. Our data are thus compatible with a COSAN assembly (aggregation number, shape, size) that remains constant from 15 to 60 -100 mM. However, they do not exclude potential macromolecular changes that could be masked by non-ideal behavior. Linear extrapolation at infinite dilution of the *s*-data above 15 mM, enables to estimate the mean *s*-value the COSAN assembly (micelles) at infinite dilution, *i*.*e*. non interacting assemblies. This gives for the micelles *s*_*micelle*_ = 0.50 +/-0.05 S in H_2_O, and *s* = 0.77 +/-0.02 S in dilute salt buffer, corresponding to *N*_agg_ = 8 and 14, respectively, in the two solvents (Figure 3d). These aggregation numbers are in the order of magnitude of those determined from isothermal calorimetry at 25°C, with *N*_agg_ = 8, increasing to 9 in 0.1 M NaCl and 14 in 1 M salt (Fernandez-Alvarez et al. 2018). Our data do not allow to confirm or infirm the pentameric state described by MD as the building block for COSAN oligomerisation. From the slope of the linear regression of *s* above the cmc, we can derive non ideality coefficients *k*_s_ of 24 +/-5 mL g^-1^ in H_2_O, and 16 +/-1 mL g^-1^ in PBS. The values of *k*_s_ depends on hydrodynamic interactions, including an obligatory repulsive contribution of excluded volume. *k*_s_ here is larger than the values of ≈ 10 mL g^-1^ measured for globular compact proteins, which have partial specific volumes similar to COSAN (Solovyova et al. 2001; Chaturvedi et al. 2018), which suggests additional electrostatic repulsions. Indeed charged COSAN micelles exhibit intermicellar repulsions of electrostatic origin (Bauduin et al. 2011). The addition of salt (or buffer) reduces these repulsions through a screening mechanism.

The number of interference fringe shifts (Δ*J*) obtained from the area under the peaks of the *c*(*s*) measures the concentration of the sedimenting material. Δ*J* is plotted as a function of concentration on Figure 4b, from data in water and diluted buffer. Analysis of SV data at 445 nm cannot be done in the same way, because *A*_445_ is saturated in the AUC above 30 mM. The general plot shape corresponds to what is typically observed for detergent micelle (Salvay and Ebel 2006; Polidori et al. 2016). Below the cmc, monomer are improperly quantified, since hardly distinguished from noise. Above the cmc, the number of interference fringes obtained from the area under the peaks of the *c*(*s*) distribution is related to the micelle concentration. It increases linearly with concentration. The refractive index increment ∂*n*/∂*c* = 0.423 +/-0.005 mL g^-1^ Δ*J* (Δ*J*/COSAN of 2.24 +/-0.03 mM^-1^ cm^-1^) is derived from slope of the linear regression (dotted line in figure 4b) of Δ*J* vs COSAN. An estimate of the cmc of 16 mM is obtained from the intercept with the concentration axis of the same linear regression. It agrees with the value of 18.6 mM determined from SANS (Bauduin et al. 2011), while being larger than that of 10 mM determined from NMR and surface tension (Gassin et al. 2015; Uchman et al. 2015). AUC, as SAXS or SANS, derives cmc as the concentration above which concentrations of monomer or premicellar species are constant, and frequently provides cmc values larger than that from techniques (surface tension, isothermal calorimetry (ITC)) measuring interactions arising in the premicellar to micellar concentration range (as examples (Salvay and Ebel 2006; Polidori et al. 2016)). The precision of our data here does not allow to evidence the effect, suggested in Figure 4a, at the lowest COSAN concentrations, as mentioned above, of solvent salt on the cmc.

To resume, COSAN is described in AUC as a monomer at up to 2-5 mM COSAN in PBS, and ≈ 10 mM in water. The monomer at larger concentrations start to self-assemble into micelles. SV-AUC is compatible, in the 15 - 100 mM range, with assemblies of COSAN with small numbers of aggregation, 8 in water, and 14 in PBS (Bauduin et al. 2011; Uchman et al. 2015; Medoš et al. 2022). This values are determined using the values of the partial specific volume of COSAN determined in water. AUC estimates the cmc of COSAN in H_2_O as in PBS, at 16 mM. AUC is unable to detect very large aggregates, such as vesicles, in the low concentration range (below the CMC). If vesicles exist in the samples, they are present in amounts too low (negligible for AUC detection), as previously concluded by SANS (Bauduin et al. 2011). The extinction coefficient at 445 nm and ∂*n*/∂*c* have been measured (Table 1), which could be used in future for studying the interaction of COSAN with other macromolecules.

### COSAN and myoglobin

We investigated myoglobin at 0.1 mM in PBS at pH 10 with COSAN at different concentrations up to 4 mM, corresponding to a ratio COSAN/myoglobin of 40. Data were acquired in the AUC at three wavelengths corresponding to absorbance maxima of myoglobin (280 and 410 nm) and COSAN (280 and 445 nm), along with interference optics. Figure 5 shows the observation of selected raw sedimentation velocity profiles of myoglobin with or without COSAN. Adding COSAN leads to a fast sedimentation of the species.

**Fig. 5.**
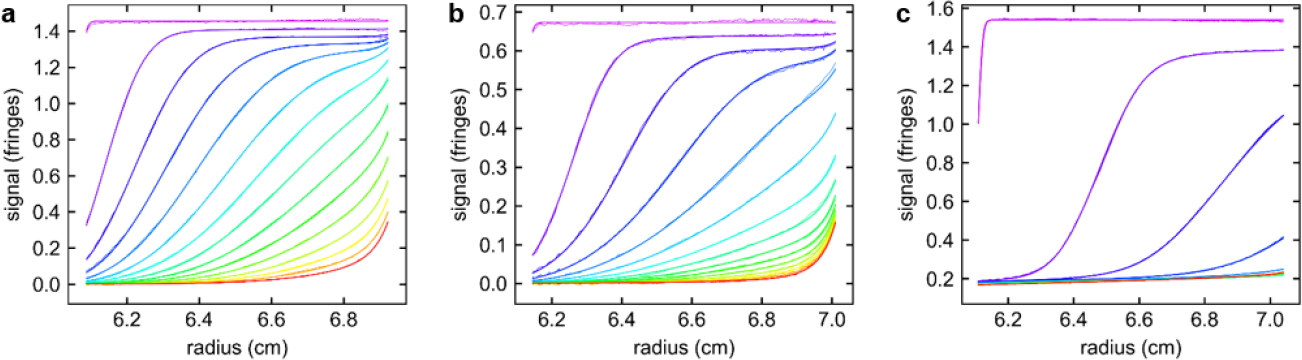
Sedimentation velocity profiles of myoglobin, in the presence of COSAN. **a**: Myoglobin alone at 0.1 mM; **b**: with 0.5 mM COSAN; **c**: with 4 mM COSAN. Data were obtained in PBS, at 20°C, at 42000 revs per min (130000 *g*), using interference optics, with optical path of 0.3, 0.15, and 0.15 cm, respectively. For clarity every third acquired scans are plotted. Time between the plotted scans is ≈ 1 h. Each panel shows the superposition of the experimental profiles -corrected by systematic noise contribution- and fitted profiles with the *c*(*s*) analysis.

Figure 6 presents, for four individual samples -COSAN alone, myoglobin alone, and mixtures at COSAN/myoglobin ratios of 1 and 5-, the *c*(*s*) obtained with various detections. At larger COSAN/myoglobin ratios, absorbance signals are saturated. COSAN at 0.1 mM can be analyzed only at 280 nm only, since it presents a very low signal with the other detections: interference and absorbance at 410 and 445 nm. It sediments very slowly, in line with the monomer state described above, at 0.36 S (Figure 6a). Free COSAN contribution is hardly discriminated from baseline in the mixtures with myoglobin, and cannot be analyzed quantitatively. Myoglobin alone sediments mainly at 1.78 +/ 0.03 S (*s*_20, w_ = 1.85 S, comparable to the published value of 1.88 S (Simmons et al. 2003)), for ≈ 95 % of the total signal (Figure 6b). The NIS analysis provides a molar mass of 17.2 kDa, close to the theoretical value of 17.4 kDa for the monomer (Chazapi et al. 2024). We used the Svedberg equation to derive a hydrodynamic radius of 2.1 nm, close to the reported value of 1.9 nm (La Verde et al. 2017), and a frictional ratio *f*/*f*_0_ of 1.21 corresponding to a globular compact shape, as expected. A minor contribution (≈ 5 %) sediments at ≈ 3 S, which could correspond to dimers.

**Fig. 6.**
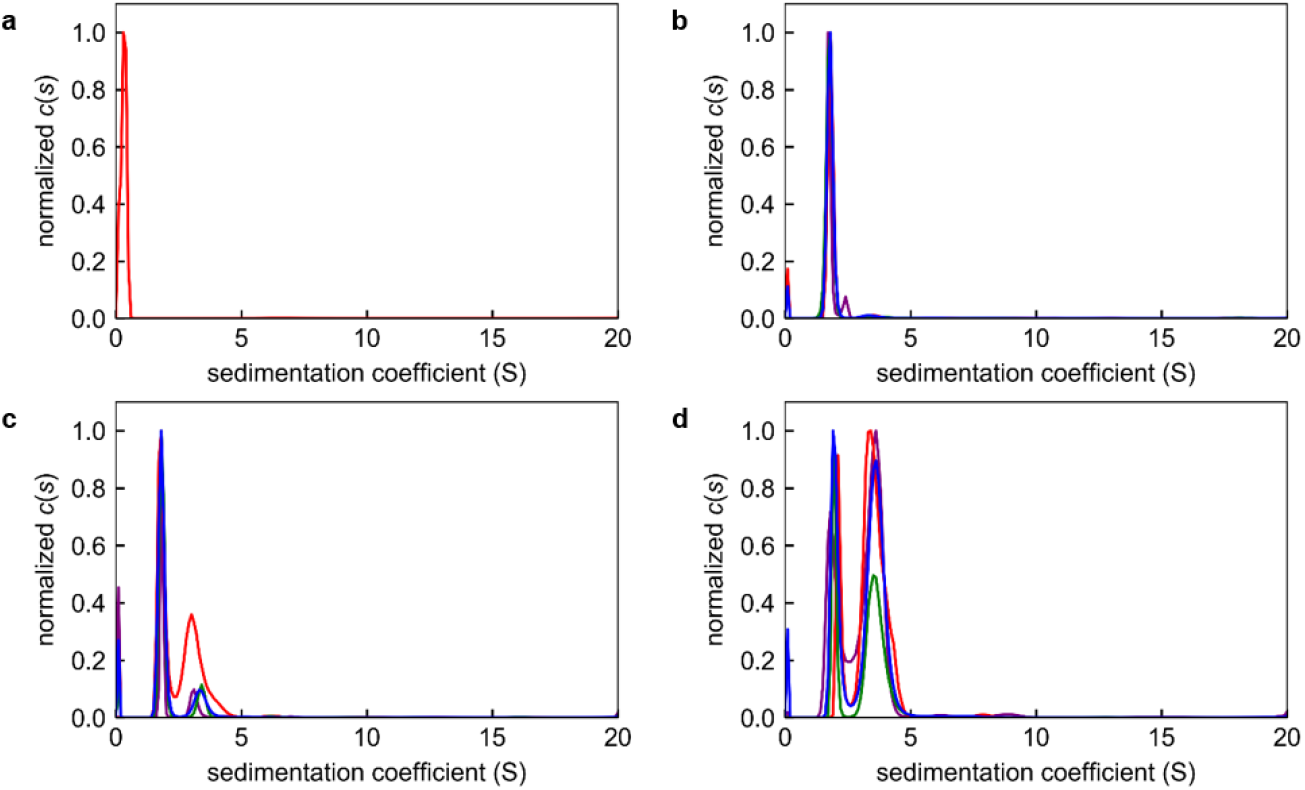
*c*(*s*) analysis of 0.1 mM COSAN, 0.1 mM myoblobin, and mixtures at ratio COSAN/myoglobin of 1 and 5. Normalized *c*(*s*), for **a**: 0.1 mM COSAN, **b**: 0.1 mM myoglobin, **c**: 0.1 mM COSAN + 0.1 mM Myoglobin, **d**: 0.1 mM COSAN + 0.5 mM Myoglobin. The *c*(*s*) were obtained with *A*_280_, *A*_445_, *A*_410_ and fringe shift detections: red, blue, green, and purple lines, respectively.

For the mixture of 0.1 mM myoglobin and 0.1 mM COSAN (Figure 6c), we observe monomeric myoglobin at 1.83 +/ 0.01 S, 17 +/-1 kDa from NIS analysis, for 50 - 80 % of the total signal. An additional contribution appears at 3.2 - 3.4 S, for 13 - 47 % of the total signal. The NIS analysis, which could be inappropriate in case of rapid equilibrium, provides an apparent molar mass of 35 - 40 kDa, close to that of the dimer. When increasing COSAN to 0.5 mM (ratio COSAN/myoglobin of 5), (Figure 6d), the contribution of the myoglobin monomer is shifted to 2.0 +/-0.1 S, and its proportion is clearly decreased (18 - 34 % of the total signal). The NIS analysis gives an apparent molar mass of 18.5 +/-1.5 kDa, slightly increased, as is the *s*-value, compared to myoglobin with no COSAN in solution, suggesting the association of one or more COSAN anions to a globular monomer of myoglobin. The main contribution of this sample (64 - 81 % of the total signal) is at 3.5 +/-0.1 S, where the NIS analysis gives an apparent molar mass of 42 +/-2 kDa, corresponding reasonably to a dimer of myoglobin with eventually bound COSAN.

The superposition of the *c*(*s*) obtained with interference optics for myoglobin and all mixtures is shown on Figure7a. The mean sedimentation coefficient above 1 S, corresponding to myoglobin assemblies, drastically increases when increasing COSAN/myoglobin ratio up to 20, and more moderately at 40 (Figure 7b). The fringe shift signal of the myoglobin assemblies decreases by ≈ 30 % when adding COSAN at ratio 1 (Figure 7c). It should remain unchanged (see the predicted trend as dashed line in Figure 7c) or slightly increase if myoglobin binds COSAN (the dotted line in Figure 7c represents the scenario where all available COSAN is bound to myoglobin). The observed signal loss indicates that myoglobin forms very large aggregates (s ≥ 25 S, M ≥ 800 kDa), which are pelleted before the first SV profiles are recorded. Above COSAN/myoglobin ratio of 1, the fringe shift signal increases regularly, and is intermediate between that calculated for 0.1 mM myoglobin alone and for 0.1 mM myoglobin with COSAN bound at its maximal value. We find plausible that what has precipitated remains precipitated and that an increasing amount of COSAN binds to the remaining ≈ 0.07 mM myoglobin. This hypothesis is supported by the fact that, beyond the first measurement after COSAN is added, the evolution of experimental Δ*J* follows the model in which all COSAN is bound (dotted line in Figure 7c). However we don’t have further argument that support this hypotheses.

**Fig. 7.**
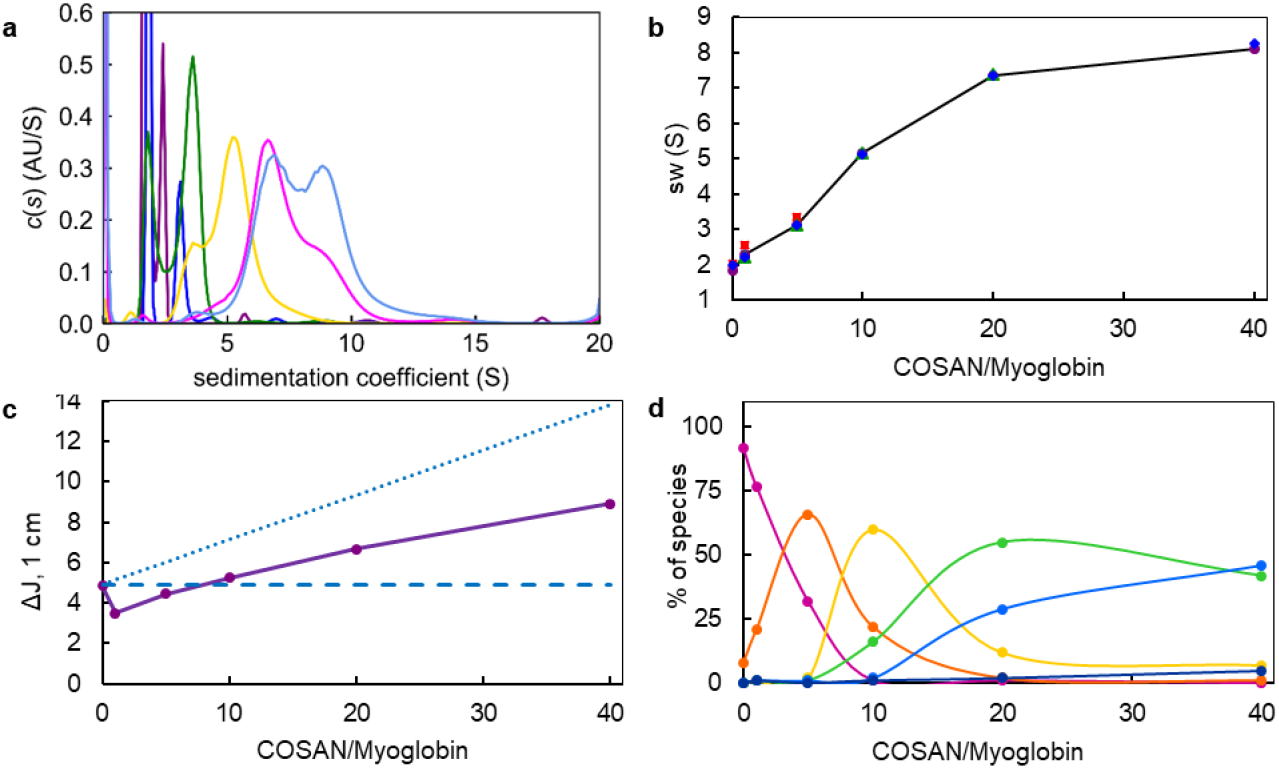
*c*(*s*) analysis of myoblobin - COSAN mixtures. **a**: *c*(*s*) obtained with interference optics, for 0.1 mM myoglobin, alone: purple; and with COSAN, at COSAN/myoglobin ratio of 1: blue; 5: green; 10: yellow; 20: pink, and 40: light blue. **b**: mean *s*-values in the 1 - 16 S range, obtained at 280 nm: red square, 410 nm: green triangle, 445 nm: blue diamond, and with interference optics: purple circles and continuous line in black; **c**: corresponding fringe shifts (purple), with the theoretical lines: for 0.1 mM myoglobin alone (dashed blue line) and for 0.1 mM myoglobin binding all available COSAN in solution (dotted blue line). **d** Distribution, in wt% quantified from *c*(*s*) obtained with interference optics integration (filled circles), of the different myoblobin sedimenting species in the presence of COSAN. The 1.8 - 2 S (pink), 3.2 - 3.5 S (orange), 5.3 S (yellow), 6.7 S (green), 8.9 S (blue), and >11.5 S (dark blue) contributions are assigned –see the text- as myoglobin monomer, dimer, and likely tetramer, hexamer, octamer, and (for simplification) 16-mers, respectively.

When increasing the COSAN/myoglobin ratio to 10, 20, and 40, species larger than the myoglobin monomer and dimer are detected, with *c*(*s*) extending to 10, 13, and 15 S, respectively (Figure 7a). Shoulders or peak maxima are found at 3.7 and 5.3 S for ratio 10, at 4.4, 6.6 and 8.9 S for ratio 20, and at ≈ 3.8, 6.9 et 8.9 S for ratio 40. The *c*(*s*) distributions of the two samples with COSAN/myoglobin ratio of 20 and 40 show their main peaks at the same *s*-values, suggesting these peaks do not correspond to reaction boundaries, i.e. mixtures of species in rapid association-dissociation equilibrium, but rather to individual, non-interacting species. While equilibrium may exist, the dissociation kinetics appear to be slow (< 10^-4^ s^-1^) (Zhao et al. 2011). The whole set of samples can be described as the superposition of sedimenting species, whose proportions, quantified from integrating the peaks of the *c*(*s*) distributions obtained with interference optics, are compiled in Figure 7d. In Figure 7d, for clarity, the 1.8 - 2 S, 3.2 - 3.5 S, 5.3 S, 6.7 S, 8.9 S, and >11.5 S contributions are designated as myoglobin monomer, dimer, tetramer, hexamer, octamer, and (for simplification) 16-mers, respectively. These assignments will be discussed below. The smallest myoglobin assemblies disappears in favor of the largest ones with increased COSAN concentrations. As summary (i) The monomer is the main species in the samples of 0.1 mM myoglobin, alone and with 0.1 mM COSAN (ratio COSAN/myoglobin of 1), (ii) the dimer is the main species with 0.5 mM COSAN (ratio COSAN/myoglobin of 5), (iii) the 5.3 S putative tetramer species with 1 mM COSAN(ratio COSAN/myoglobin of 10), and (iv) the 6.7 S and 8.9 S –putative hexamer and octamer-are the two main species of the samples with 2 and 4 mM COSAN (ratio COSAN/myoglobin of 20 and 40, respectively).

To decipher myoglobin - COSAN stoichiometry for each sedimenting species, we would aim to combine the signals obtained with the different optics. In the specific case of the interactions of myoglobin with COSAN, this approach founds however strong limitations, since the UV-visible spectra of myoglobin is deeply affected by COSAN (Goszczyński et al. 2017; Chazapi et al. 2024). Chapazi (Chazapi et al. 2024) *e*.*g*. describes a strong absorbance decrease (by a factor of ≈ 30 % for COSAN/myoglobin ratio of 10), with a slight red shift, of the Sorret band at 410 nm, and an absorbance increase in the 480 nm zone. Signal analysis of the SV profiles at 410 nm and 445 nm thus cannot be used to decipher stoichiometry. Interference fringe shift and absorbance at 280 nm signals, Δ*J* and *A*_280_, can thus only be used. The two optics are highly complementary, since, for a same concentration and optical path length, *A*_280_ / Δ*J* ratios are extremely different for COSAN and myoglobin, with values of 0.1 and 1.6, respectively, from Table 1. The analysis required quantitative values for the refractive index increments and extinction coefficients (see Materials and Method section). For myoglobin, Δ*J* derived from the calculated *c*(*s*) coincides perfectly (99 %) with the value expected from theoretical value whereas *A*_280_ from the *c*(*s*) is significantly larger. We will consider *A*_280_ / myoglobin determined in AUC (Table 1). For COSAN, we used ∂*n*/∂*c* = 0.42 mL g^-1^ determined above, and *A*_280_ / COSAN obtained by UV-visible spectrophotometry, from a dilution series in water of our COSAN sample.

In the sample comprising 0.1 mM myoglobin and 0.1 mM COSAN, the 1.8 S peak, from Δ*J* and *A*_280_, would comprise 55 µM of myoglobin and 8 µM de COSAN (thus 0.2 molecules of COSAN per myoglobin), and the 3.2 S peak one to 12 µM of myoglobin and 55 µM of COSAN (thus 5 molecules of COSAN per myoglobin). The total amounts detected in the AUC (67 µM of myoglobin and 62 µM of COSAN) corresponds to ≈ 65 % of the loaded (100 µM of each) amounts. Thus 35 % of the material is missing, which was potentially pelleted fast, being precipitated or forming large aggregates. We analyze also the sample with 0.1 mM myoglobin and 0.5 mM COSAN, which has an experimental absorbance at 280 nm of 1.6 absorbance unit, above but close to the limit for absorbance linearity. The 2 S peak would comprise 27 µM of myoglobin and 46 µM de COSAN (thus COSAN/myoglobin = 1.7), and the 3.5 S peak, 46 µM of myoglobin and 307 µM of COSAN (thus COSAN / myoblobin = 7). The total amounts detected in the AUC are thus 74 and 353 µM of myoglobin and COSAN, respectively, corresponding to 70-75 % of the loaded material. The calculated *s*-values for a globular compact dimer of myoglobin binding 5 or 7 COSAN are slightly smaller but in line with the experimental values.

To describe the larger aggregates, at *s* = 5.3, 6.7 and 8.9 S, only the *s*-values can be considered since *A*_280_ > 2 in the AUC. We argued above that the 6.7 and 8.9 S contributions can be considered as related to species. They are compatible with a hexamer and an octamer, respectively, of myoglobin binding 10 and 18 COSAN / monomer of myoglobin. The 5.3 S contribution was observed only in the sample with a COSAN/myoglobin ratio of 10. The *s*-value is consistent with that of tetramer binding 8 COSAN per monomer of myoglobin, but we cannot ascertain it does correspond to a mean of smaller and larger oligomers. These conclusions are compiled in Table 2.

**Table 2.**
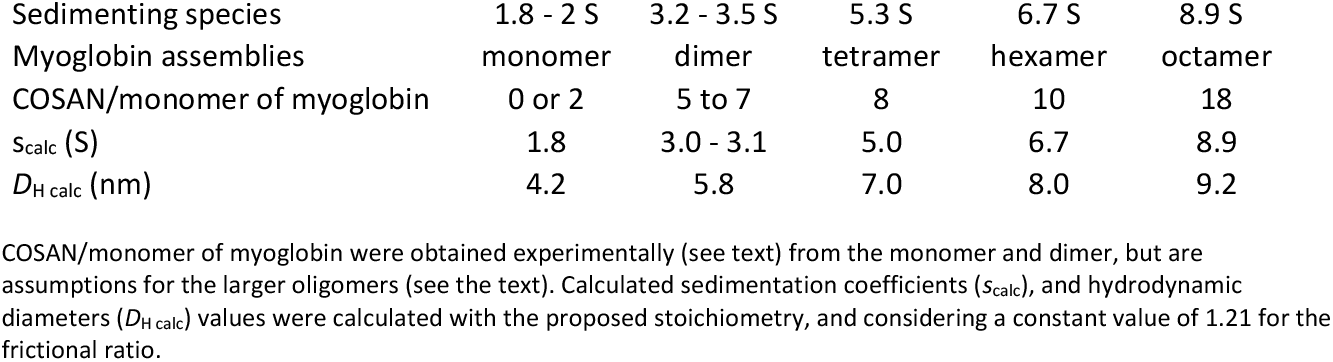
Myoglobin putative association states in the presence of COSAN.

The AUC experiments provide a description of myoglobin - COSAN samples in terms of mixtures of myoglobin oligomers. We aim to compare these results with published SAXS and dynamic light scattering (DLS) results (Chazapi et al. 2024). We present on Figure 8a the mean aggregation numbers, *N*_agg_, i.e. the aggregation number of myoglobin proteins in the assembly formed upon addition of COSAN, derived from SAXS and AUC. SAXS analysis were done with samples at the same myoglobin and COSAN concentrations as in this work, but only in water. *N*_agg_ from AUC were calculated with the relative concentration (wt %) of the proposed myoglobin association states provided in Figure 7d. We considered arbitrarily the species sedimenting above 11.5 S as 16-mers. SAXS and AUC analysis show a qualitative agreement with an increase in *N*_agg_ of the myoglobin when adding COSAN (Figure 8a). At COSAN/Myoglobin ratio of 40, *N*_agg_ from SAXS is 18 while from AUC, *N*_agg_ is 7. This difference is most probably related to the depletion in AUC of very large myoglobin aggregates, which is evidenced from fringe shift quantification in Figure 7c. Because of the experimental setting (with seven cells multiples detection), data were acquired in the AUC every 20 min for each sample/detection, which does not allow to characterize easily > 20 – 25 S species (*N*_agg_ ≈ 40), and particularly if they are in minor amounts.

**Fig. 8.**
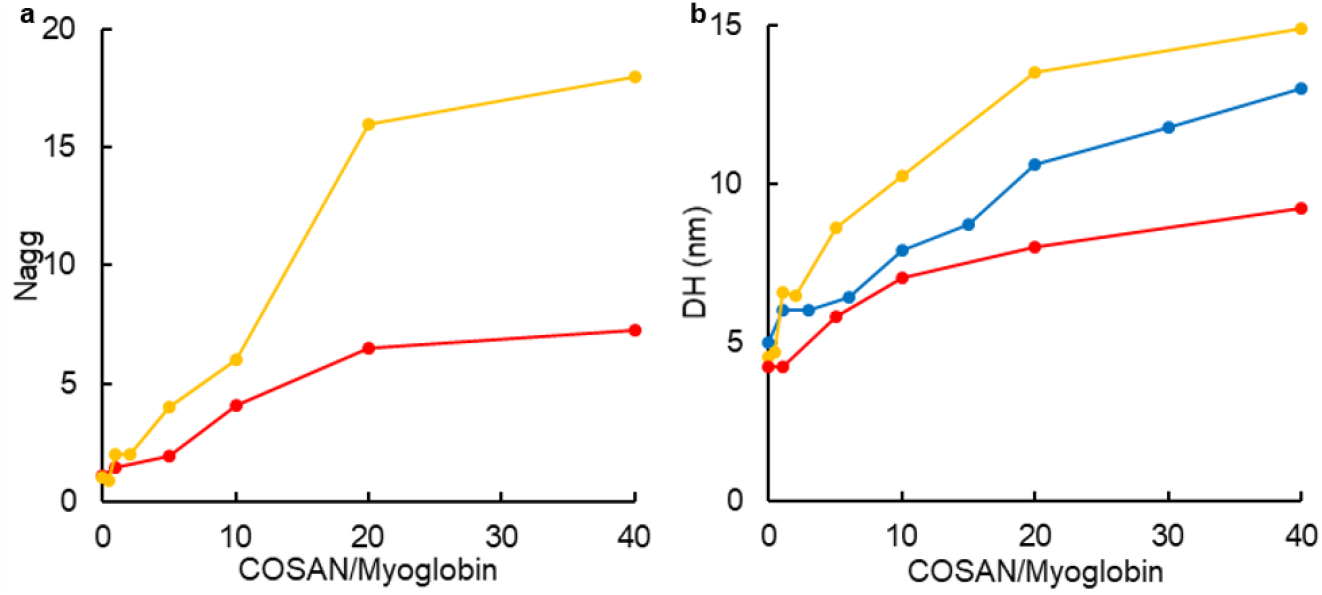
Comparison of myoglobin mean size by AUC, SAXS and DLS. **a**: Mean aggregation numbers, from AUC in red (this work) and from SAXS in yellow; **b**: Hydrodynamic diameters from DLS in blue; derived from SAXS analysis in yellow; and corresponding to the main assemblies from AUC in red. Myoglobin concentration was 0.1 mM in SAXS and AUC, and 0.01 mM in DLS. Solvent was PBS in AUC and DLS, and water in SAXS experiments. SAXS and DLS data are from Chazapi I, Merhi T, Pasquier C, Diat O, Almunia C, Bauduin P (2024) Angew Chem Int Ed Engl e202412588. doi: 10.1002/anie.202412588 (Chazapi et al. 2024).

Scattering techniques provide information of the size of the assemblies and hence can be compared to AUC results. We have reported on Table 2 the hydrodynamic diameters, *D*_H_, corresponding to the major species determined by AUC. *D*_H_ were measured using DLS for myoglobin at 0.01 mM in PBS with COSAN/myoglobin ratio reaching 40 (Chazapi et al. 2024). We also calculated the *D*_H_ values corresponding to the dimensions of the ellipse model derived from SAXS, in water, for each of the different COSAN/myoglobin ratio (Chazapi et al. 2024). As seen in Figure 8b, the derived *D*_H_ values are in agreement, given AUC values are only for the main species, while scattering techniques give mean values in which the signal of the larger species is amplified, and were obtained in different experimental conditions: solvent was PBS in AUC and DLS, and water in SAXS; concentration was 0.1 mM in SAXS and AUC, and 0.01 mM in DLS.

To summarize, SV-AUC experiments show that adding COSAN to myoglobin lead to the formation of very large aggregate, with e.g. approximatively 30% of the material pelleted at low angular velocity as characterized for COSAN/myoglobin ratio of 1 and 5. While myoglobin alone is in a monomeric state, the addition of COSAN induces oligomerization: primarly dimers at a COSAN/myoglobin ratio of 5, and likely tetramers at a ratio of 10, hexamers and octamers at the larger ratios of 20 and 40. COSAN binds to myoglobin in substantial amounts (e.g. ≈ 12 COSAN anions per dimer). If not all, a significant fraction of COSAN in solution is bound to myoglobin. The sizes of the assemblies described by AUC are in agreement with previously published SAXS and DLS results, due to specific limitations and hypotheses associated with the different techniques.

## Conclusion

This work aimed to explore the potential and limitation of AUC for characterizing COSAN assemblies and macromolecular assemblies in the presence of COSAN. We have investigated two systems, COSAN alone and myoglobin with COSAN, both of which have been studied previously using a variety of techniques, including SAXS, SANS, DLS, ITC and MD. The results for both systems are in qualitative agreement with those obtained from these other techniques. AUC effectively describes the self-assembly of COSAN above the mM range (> 10 mM in water, > 2 - 5 mM in PBS) into micelles. Small numbers of aggregation, 8 in water and 14 in PBS, are derived from AUC, which does not provide evidence for larger assemblies, such as vesicles, previously observed in other investigations. The fact that COSAN assemblies sediment slowly (< 1 S) is advantageous for studying the effect of COSAN on the assemblies of macromolecules that sediment at faster rates using SV-AUC. AUC clearly shows that myoglobin self-assembles into small multimers in the presence of COSAN. By superimposing the *c*(*s*) profiles obtained at different COSAN/myoglobin ratios, while keeping myoglobin concentration constant, the formation of various assemblies, up to octamers, is suggested. A significant proportion (≈ 30%) of myoglobin forms large aggregates, –which are detected by their absence in AUC measurement. The determination of the amount of bound COSAN, based on combining signals from different detection methods, was limited by wavelength, as the binding process affects the spectroscopic properties of both myoglobin and COSAN at 410 and 445 nm. The very strong absorbance of COSAN at 280 nm allows for this estimation, but it is limited to COSAN concentration below 0.5 mM. Our estimates, however, shows that a significant proportion of the COSAN in solution is bound to myoglobin. The fact that SV-AUC can relatively easily discriminate between different protein association states makes it a valuable tool for investigating protein interactions mediated by COSAN. Combining different detection methods in AUC enables the analysis of bound COSAN for each association state, which, despite the limitations observed here for myoglobin, offers promising perspectives for understanding the link between macromolecule-macromolecule and COSAN-macromolecule interactions.

## Materials and Methods

### Materials

The 3,3’-commo-bis[closo-1,2-dicarba-3-cobalta dodecaborane] anion was purchased from Katchem in cesium form (Cs[(C_2_B_9_H_11_)_2_Co], COSAN, > 98%). We used diethyl ether ((CH_3_CH_2_)_2_O, ≥ 99.7%) and sodium chloride (NaCl, ≥ 99%) from Sigma Aldrich for the ion exchange to produce the sodium salt of COSAN from the cesium salt. The sodium salt of COSAN was used in this study. The counter-ion exchange of COSAN was performed following the procedure described by Merhi et al (Merhi et al. 2020). Myoglobin from equine skeletal muscle was purchased from Sigma Aldrich.

### AUC-SV theoretical background

Sedimentation profiles, reflecting the evolution of concentration, *c*, with time, *t*, and radial position, *r*, are monitored in real time using optical systems in the analytical ultracentrifuge. Analysis are based on numerical solutions of the Lamm equation, describing rigorously the transport of each type of macromolecule in the ultracentrifuge, as a function of their sedimentation and diffusion coefficients, *s*, and *D*, respectively:

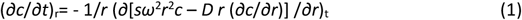

*c* is the concentration, *ω* the angular velocity. The Lamm equation can be used to analyse complex systems, e.g. mixtures of non-interacting or interacting molecules. The Svedberg equation relates, for an ideal two-component solution, *s* and *D* to the buoyant molar mass 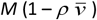, with *M* and *v* the molar mass and partial specific volume of the molecule, and *ρ* the solvent density:

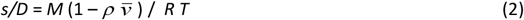

where *R* is the universal gas constant and *T* the temperature in Kelvin. The Svedberg equation, combined with the Stokes–Einstein relationship, linking *D* to the hydrodynamic diameter, *D*_H_, can be written

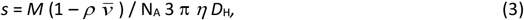

with N_A_ Avogadro’s number and *η* the solvent viscosity.The sedimentation coefficient can be expressed as *s*_20,w_, after correction for solvent density and viscosity to the solvent density,ρ_20,w_, and viscosity, *η*_20,w_, of water at 20°C:

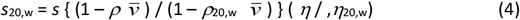

*D*_H_ is related to *D*_0_, the diameter of the anhydrous volume, by the frictional ratio *f*/*f*_0_ *f*/*f*_0_ = *D*_H_ /*D*_0_, (5)

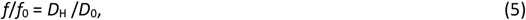

*f*/*f*_0_ depends on particle shape and hydration. The effect of shape for compact assemblies are moderate. We typically consider *f*/*f*_0_ = 1.25 for globular compact macromolecules. We found here *f*/*f*_0_ = 1.21 for myoglobin. *f*/*f*_0_ changes from 1 for an anhydrous sphere to 1.2 or 1.5 for anhydrous ellipsoids of axial ratio 4 or 10. For unfolded proteins, *f*/*f*_0_ depends on the molar mass and reaches 3 for *M* = 300 kDa (Salvay et al. 2012).

COSAN in salted buffer is a three-component system, as composed of water (component 1), the COSAN (composed of an anion and Na^+^, component 2), and solvent salt (NaCl, component 3). The buoyant mass is *M* (∂*ρ*/∂*c*)_µ_, (∂*ρ*/∂*c*)_µ_, being the density increment of COSAN at constant chemical potential of water and solvent salt. The Svedberg equation, in the limit of infinite dilution, may be written:

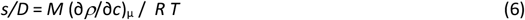

(∂*ρ*/∂*c*)_µ_ can be written in terms of a preferential solvent interaction parameters, (∂*w*_3_/∂*w*_2_)_µ_ in g/g, or (∂*m*_3_/∂*m*_2_)_µ_ in mol/mol, respectively, which expresses the amount of salt (component3) that would have to be added to (or removed from) the solvent system to restore thermodynamic equilibrium when COSAN is added (for reviews, see (Timasheff 1993; Eisenberg 2000; Ebel 2016).

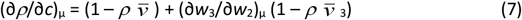

Full dissociation of Na^+^ from COSAN would induce solvent rearrangements corresponding to (∂*m*_3_/∂*m*_2_)_µ_ = -0.5 mol mol^-1^, (∂*w*_3_/∂*w*_2_)_µ_ = (*M*/*M*_3_)(∂*m*_3_/∂*m*_2_)_µ_ = -0.084 g g^-1^. We can calculate *v* = 0.736 mL g^-1^ at 20°C in water, from reported densities of COSAN solutions at up to 1 M COSAN in water (Medoš et al. 2022), and *v* _3_ = 0.3 mL g^-1^ in 0.15 M NaCl from tabulated density tables. Using these values and a solvent density of 1.005 g mL^-1^, (∂*ρ*/∂*c*)_µ_ for fully dissociated COSAN would be 0.201, to be compared to 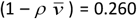. The molar mass of COSAN assemblies would be 30% larger than that calculated without considering ion dissociation and with 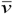 measured in water. COSAN is however described with negligible ion binding in pure water (Medoš et al. 2022). We thus decided to keep the experimental value determined in water 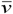 of 0.736 mL g^-1^ for our analysis.

In non-ideal solutions, the macromolecule concentration influences *s* and *D* through spatial and velocity correlations between particles. The SV profiles are significantly affected by the concentration dependency of *s*, and weakly by that of *D* (Solovyova et al. 2001; Chaturvedi and Schuck 2024). For repulsive weak interactions (i.e. excluded volume and electrostatic interactions), *s* decreases when increasing concentrations, and sharpening of the boundary is observed. For moderate protein concentrations, *s* can be described in the linear approximation:

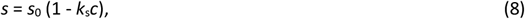

where *s*_0_ is the extrapolated to infinite dilution sedimentation coefficients, and *k*_s_ the related concentration dependency parameter.

### AUC-SV principle of analysis

The continuous distribution analysis *c*(*s*): The measured sedimentation signal is considered as a superposition of independently sedimenting species, each described by a Lamm equation solution. The sample is described as a mixture of typically 200 non-interacting particles, defined by their minimum and maximum *s*-values. These particles are assumed to have the same 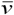 and *f*/*f*_0_ values (that can be fitted), allowing to fix a relationship between *s* and *D*. This, and efficient noise decomposition techniques, allow to obtain diffusion deconvoluted sedimentation coefficient distributions *c*(*s*) (Schuck 2000) (Schuck 2000). Integration of the *c*(*s*) peaks provides values for the mean sedimentation coefficient and the signal related to species concentration. The *c*(*s*) peaks may represent non-interacting species, or reaction boundaries - i.e. mixtures of species in a dynamic equilibrium of association dissociation (Zhao et al. 2011) . When peaks superpose in the *c*(*s*) of samples at different concentrations, we consider they represent non-interacting species. The values of *s* can then be analysed using the Svedberg equation (Equation 3). The non-interacting species (NIS) analysis: for homogeneous samples, it is possible to use the Lamm equation to fit *s* and *D*, and thus determine without hypothesis 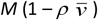 and *D*_H_. The NIS analysis can be extended to relatively homogeneous samples, considering *e*.*g*. two or three species in solution. The NIS analysis is not adapted to macromolecular systems undergoing equilibrium of association, unless the kinetics of dissociation is low (Zhao et al. 2011) . These analyses are available in two free to download packages: SEDFIT, which we use here, and UltraScan (https://www.ultrascan3.aucsolutions.com/).

For multicomponent species, signal analysis allows getting information on stoichiometry. Absorbance signal, *A*, is related to concentration by the Beer–Lambert law and appropriate extinction coefficient; fringe shift signal, Δ*J*, obtained when using interference optics in the AUC, is related to concentration, *c* (g/mL), optical path length, *l* (cm), increment of index of refraction, ∂*n*/∂*c* (ml/g), of the particle, and laser wavelength (*λ* = 6.75 10^-5^cm in our instrument):

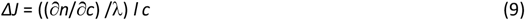

The composition of a complex comprising two different components behaving optically differently may be determined combining two signals from two different optics, such as interference and absorbance, or absorbance at two different wavelengths.

### AUC-SV samples, experiments and analysis

A carefully prepared aqueous stock solution of COSAN at 263.45 mM was used to prepare, by dilution, the samples in H_2_O, or 20 mM Tris-HCl pH 8, 150 mM NaCl, or PBS pH 7.4, at various concentrations ranging from 2 to 100 mM. Myoglobin samples at 0.1 mM were prepared in PBS buffer pH 10, without and with COSAN at various concentrations up to 4 mM. AUC-SV experiments were conducted in an XLI analytical ultracentrifuge (Beckman, Palo Alto, CA) using ANTi-50 rotor, using double channel Titanium centerpieces (Nanolytics, Germany) of typically 1.5 mm optical path length equipped with sapphire windows. Acquisitions were done at 42,000 revs per min (130,000 *g*), at 20°C overnight, using absorbance detection at 280, 410 and 445 nm and interference detection. Data processing and analysis were done using the program SEDFIT version 16.36 and 17 (Schuck 2000) from P. Schuck (NIH, USA), free available at https://sedfitsedphat.nibib.nih.gov/software. REDATE version 1.0.1 (Zhao et al. 2015) and GUSSI version 1.4.2 and 2.1.6 (Brautigam 2015) from C. Brautigam (USA), free available at https://www.utsouthwestern.edu/research/core-facilities/mbr/software/. The program SEDNTERP 3.0.2beta (Philo 2023), free available at http://www.jphilo.mailway.com/, was used for calculation of buffer density, *ρ*, and viscosity, *η*. For water, *ρ* = 0.99828 g mL^-1^, *η* = 1.002 cp; for PBS, *ρ* = 1.006 g mL^-1^, and *η* = 1.02 cp. The partial specific volume 0.744 mL g^-1^ and the refractive index increment (∂n/∂c) of myoglobin of 0.188 mL g^-1^ of myoglobin were calculated from amino-acid composition with SEDFIT.

### Extinction coefficients of COSAN

From wavelength scans at 3000 revs. per min, of COSAN, at 15 mM and 30 mM in PBS, and at 30 mM in H_2_O, we derive *A*_445_ = 0.33 mM^-1^ cm^-1^, which was used for further analysis in that work. We checked, for COSAN samples at 2, 5, 10 and 15 mM, in water and / or PBS, prepared with precise weighted dilutions, that the absorbance at 445 nm was the same when measured in a spectrophotometer (Nanodrop 2000) and in the AUC running at low angular velocity (with a wavelength scan at 3000 revs per min, 665 *g*, for a radial position of 6.5 cm corresponding to the central position of the channel). The measurements were repeated after one day with the spectrophotometer, and 6 days in AUC, to probe eventual time after dissolution effects. Because COSAN absorbs hugely at 280nm, we checked also the same signal at 445 nm was obtained in the AUC with or without a cut-off filter below 400 nm. UV light contaminating light in the visible indeed may arise from higher order reflections from the diffraction grating monochromator (Schuck, P., Zhao, H., Brautigam, C. A., & Ghirlando, R 2016). There are no significant differences between nanodrop and AUC measurements, with or without optical filter. We obtained, in H_2_O: *A*_445_ = 0.352 +/-0.005 mM^-1^ cm^-1^, and in PBS: *A*_445_ = 0.334 +/-0.006 mM^-1^ cm^-1^, which are not significantly different. These values are comparable to reported values (0.443 (Fink et al. 2023; Chazapi et al. 2024); 0.335 - 0.555 estimated spectroscopically from different COSAN solutions over the years by us). We determined from wavelength scans *A*_410_ = 0.21 +/-0.01 mM^-1^ cm^-1^, in PBS and H_2_O. We used *A*_280_ = 24 mM^-1^ cm^-1^, determined by us in water and PBS with a spectrophotometer. This value is smaller but comparable to reported values of 26.5 (Chazapi et al. 2024), or 30.9 (Fink et al. 2023).

### Calculation of D_H_ for ellipsoids

SAXS analysis leaded to myoglobin complexes described as triaxial ellipsoids (Chazapi et al. 2024). We calculated *D*_H_ of derived biaxial ellipsoids -considering for the smallest axis of the biaxial ellipsoid the mean values of the two smaller dimensions of the triaxial ellipsoid from SAXS-using the program ellipscylin-10 (García De La Torre and Hernández Cifre 2020), free available at http://leonardo.inf.um.es/macromol..

## Acknowledgments

This work used the platforms of the Grenoble Instruct-ERIC center (ISBG ; UAR 3518 CNRS-CEA-UGA-EMBL) within the Grenoble Partnership for Structural Biology (PSB), supported by FRISBI (ANR-10-INBS-0005-02) and GRAL, financed within the University Grenoble Alpes graduate school (Ecoles Universitaires de Recherche) CBH-EUR-GS (ANR-17-EURE-0003). This work was funded by the Agence Nationale de la Recherche (ANR-22-CE06-0026 Promenix). The authors thank Alban Jonchère for his support in spectroscopy.

